# Signaling by Ras^G12V^ depends on EGFR activity *in vivo*

**DOI:** 10.1101/2025.11.12.687982

**Authors:** Patricia Vega-Cuesta, Ana López-Varea, Ana Ruiz-Gómez, Jose F. de Celis

## Abstract

Ras proteins are key regulators of signaling pathways initiated by transmembrane tyrosine kinase receptors, and activating Ras mutations are drivers in cancer and RASophaties. The development of effective therapies against these diseases requires a profound understanding of how Ras mutations modify pathway output and interact with other pathway components. In this work we generated G12V activating mutations in the *Drosophila Ras1* locus and used *Ras1^G12V^* and human *KRASB^G12V^* to analyze *in vivo* the consequences of endogenous Ras^G12V^ expression and its interplay with other components of the EGFR/Raf/MAPK pathway. We find that Ras^G12V^ alters the timing but not the spatial pattern of ERK activation. Consistently, Ras^G12V^ mutant proteins respond to EGFR stimulation and can partially compensates for EGFR loss to generate basal levels of ERK activity. However, full pathway activation is still strictly dependent on EGFR function. Altogether, our results suggest that Ras^G12V^ mutant proteins act as an amplifier of EGFR stimulation accelerating and raising baseline ERK activity but requiring EGFR signaling for maximal activation. This mechanistic framework supports the use of combined inhibition of both Ras and EGFR to maximize therapeutic efficacy in Ras^G12V^ tumors.

## Introduction

Ras GTPases are members of signaling pathways that play fundamental roles in the control of cell proliferation and differentiation during multicellular development and tissue homeostasis (Tidyman & Rauen, 2009; Simanshu *et al*, 2017). They are among the most common alterations in human cancers and are also linked to a group of hereditary diseases named RASophaties (Tidyman & Rauen, 2009; Hebron *et al*, 2022). Ras oncogenic mutations typically occur at one of three hotspots, Gly12, Gly13, or Gln61, leading to the accumulation of Ras in its active, GTP-bound form (Scharpf *et al*, 2022; Prior *et al*, 2020). Specific amino acid substitutions in these positions produce different effects in the Ras activation/deactivation cycle, protein stability and/or interactions with downstream effectors, thereby generating variable levels of Ras activity (Muñoz-Maldonado *et al*, 2019). For example, amino acid changes in G12, the most frequently mutated residue in human tumors, impair GAP-mediated GTP hydrolysis while preserving guanine nucleotide exchange and GEF interactions, resulting in a moderately activated Ras protein (Muñoz-Maldonado *et al*, 2019).

Early studies suggested that oncogenic Ras proteins drive growth factor–independent activation of the Raf/ERK pathway and were therefore assumed to function as constitutively active forms of Ras proto-oncogenes (Rajalingam *et al*, 2007). This view was consistent with the observed resistance of Ras-mutant tumors to EGFR-targeted therapies (Douillard *et al*, 2013; Lièvre *et al*, 2006; Amado *et al*, 2008). More recently, however, dual inhibition of EGFR and downstream Ras signaling components has emerged as a promising therapeutic strategy in RAS-mutant cancers (Tatli & Doganay, 2021). Moreover, studies in mouse models have shown that EGFR is required for Ras-driven pancreatic and lung tumorigenesis (Navas *et al*, 2012; Moll *et al*, 2018), and mutated Ras proteins respond to receptor stimulation *in vitro* (Hood *et al*, 2019). Thus, the degree to which Ras oncogenes require upstream activation for signaling, and the role of EGFR in Ras-driven pathogenesis, remain unresolved questions.

Unravelling the complex biology of mutant Ras isoforms demands experimental models allowing the evaluation of specific Ras mutations in controlled genetic backgrounds. One organism that fulfills these requirements is *Drosophila melanogaster*, as it contains a single Ras ortholog to human *K/N/H-RAS* genes, named *Ras1*. Furthermore, the set of Ras upstream regulators and downstream effectors display widespread conservation and low gene redundancy (Mirzoyan *et al*, 2019). This has facilitated the analysis of the functional requirements of the Ras pathways during normal development, as well as the identification of additional modulators of Ras signaling (Karim *et al*, 1996; Huang & Rubin, 2000). However, previous studies have relied on ectopic expression of mutated Ras using the Gal4/UAS system (mostly G12V) (Brumby & Richardson, 2003; Pagliarini & Xu, 2003) a genetic background that dramatically alters the ratio between mutated versus normal Ras proteins and may change the genomic context in which Ras mutations drive tumorigenesis.

In this work, we developed genetically modified *Drosophila* strains in which either activated versions of *Drosophila Ras1* or human *KRASB* are expressed from the endogenous *Ras1* locus. We focused on the G12V mutation, which is frequently found in human cancers (Prior *et al*, 2020) yet lack effective clinical inhibitors (Kwan *et al*, 2022). We used the wing imaginal disc to define the timing and extent of ERK activation *in vivo* and investigated the interactions of Ras^G12V^ with upstream and downstream components of the EGFR pathway. We find that endogenous expression of *Ras^G12V^*increases basal ERK activity but preserves the normal spatial pattern of EGFR activation. Notably, Ras^G12V^ mutations remain sensitive to EGFR receptor activation and require EGFR function to achieve high level pathway activity, and consequently they cannot be considered as fully constitutive. These results align with the observed dependence of Ras1^G12V^ on growth factors (Hood *et al*, 2019), and support therapeutic strategies that combine EGFR inhibition with downstream blockade of Ras signaling in tumors harboring Ras mutations.

## Results

### Generation and genetic characterization of *Ras1^KO-Kin^* transgenic strains

We replaced the *Ras1* coding region with an *attP* cassette by homologous recombination at the 5’ and 3’ ends of *Ras1* gene. The resulting allele (*Ras1 knockout*; *Ras1^KO^*) behaves as a *Ras* null allele and fails to complement hypomorphic *Ras* alleles or a *Ras* deficiency (not shown). Using this *attP* landing site, we reintroduced different wild type or activated forms of *Drosophila Ras1* and human *KRASB*, generating a collection of *Ras1 knock-in (Ras1^Kin^*) strains (Fig. 1A).

**Figure 1.**
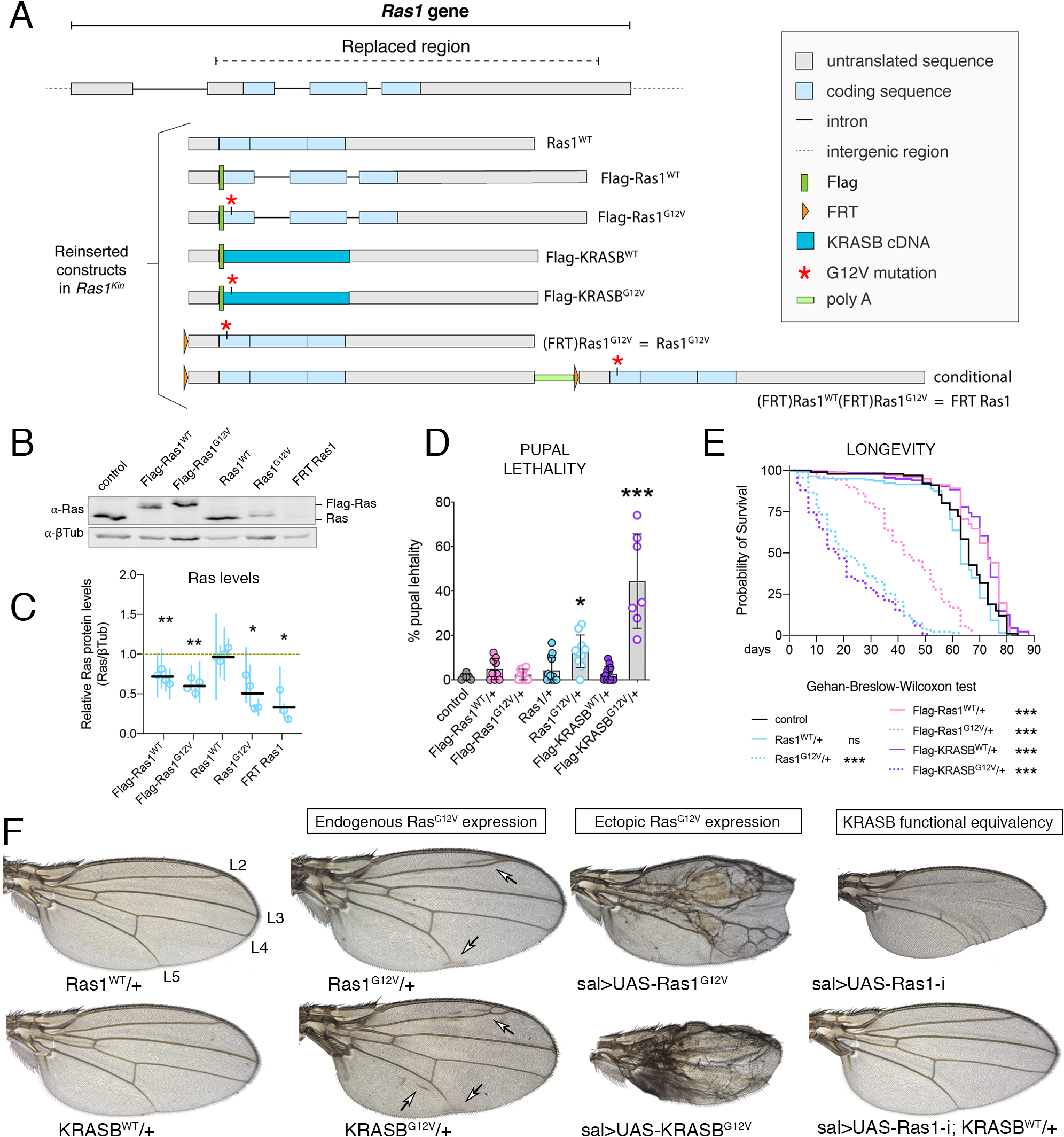
Ras1^KO-Kin^ constructs and adult phenotypes. (A) Schematic representation of the *Ras1* genomic region showing the extent of the *Ras1^KO^* deletion (“replacer region”; up) and the molecular structure of *Ras1^KO-Kin^* constructs. From top to bottom: *Ras1^WT^*, *Flag-Ras1^WT^*, *Flag-Ras1^G12V^*, *Flag-KRASB^WT^*, *Flag-KRASB^G12V^*, *FRT Ras1^G12V^*(referred as *Ras1^G12V^*) and *FRT Ras1^wt^ FRT Flag-Ras1^G12V^* (referred as *FRT Ras1*). (B) Representative western blot from third instar larval extracts of the following genotypes hemizygous for *Ras1*: *Df(3R)by10/+* (control), *Flag-Ras1^wt^/Df(3R)by10*, *Flag-Ras1^G12V^/Df(3R)by10*, *Ras1^wt^/Df(3R)by10*, *Ras1^G12V^/Df(3R)by10* and *FRT Ras1/Df(3R)by10*. Note the differential migration for tagged (Flag-Ras; up) versus untagged (Ras; down) proteins. βTub was used as an internal loading control. (C) Quantification of Ras protein levels relative to βTub corresponding to three biological replicates of the western blots shown in B. Horizontal black lines indicate the mean of biological replicates. Each vertical blue line indicates the standard deviation (SD) of technical replicates, and the circles represents the mean of each biological replicate. (D) Percentage of pupal lethality in control and *Ras1^KO-Kin^*heterozygous backgrounds. The bar graphs display the mean ± SD. (E) Survival curves of heterozygous females for control, *Ras1^WT^/+* (blue line)*, Ras1^G12V^/+* (dotted blue line)*, Flag-Ras1^WT^* (pink line)*, Flag-Ras1^G12V^ /+* (dotted pink line)*, Flag-KRASB^WT^/+* (purple line)*, and Flag-KRASB^G12V^/+* (dotted purple line) heterozygous females. Survival statistic were calculated using Gehan-Breslow-Wilcoxon test. (F) Adult female wing phenotypes of heterozygous *Ras1^WT^* and *KRASB^WT^*(left), heterozygous *Ras1^G12V^* and *KRASB^G12V^*(“Endogenous Ras^G12V^ expression”), *sal^EPv^-Gal4 UAS-GFP/UAS-Ras1^G12V^* (sal>UAS-Ras1^G12V^) and *sal^EPv^-Gal4 UAS-GFP/UAS-KRASB^G12V^* (sal>UAS-KRASB^G12V^; “Ectopic Ras^G12V^ expression”), and functional rescue of *Ras1* knockdown (*sal^EPv^-Gal4 UAS-GFP/UAS-Ras1-RNAi*; sal>UAS-Ras1-i) by endogenous *KRASB^WT^* expression in *sal^EPv^-Gal4 UAS-GFP/UAS-Ras1-RNAi; KRASB^WT^/+* flies (sal>UAS-Ras1-i; KRASB^WT^/+; “KRASB functional equivalence”).

Quantification of Ras protein levels from hemizygous larval extracts revealed that untagged wild type *Ras1* strains produce normal Ras protein levels, whereas tagged (*Flag-Ras1^WT^*) and conditional (*FRT Ras1*) lines have reduced amounts of Ras protein (71% and 34% compared to control, respectively; Fig. 1B-C). Despite repeated attempts, we were unable to obtain transgenic flies for the untagged *Ras1^G12V^* constructs. Instead, untagged *Ras1^G12V^*was generated from the *FRT Ras1* conditional expression line (*(FRT)Ras1^G12V^*, hereafter referred to as *Ras1^G12V^*; Fig. 1A). G12V mutant lines (*Flag-Ras1^G12V^*and *Ras1^G12V^*) display reduced Ras protein levels compared to controls (65% and 50%, respectively; Fig. 1B-C).

Heterozygous flies bearing activated forms of Ras1 or KRASB were viable but cause variable levels of dominant lethality (Fig. 1D). The surviving heterozygous adults displayed reduced longevity (Fig. 1E), and phenotypes indicative of excess Ras signaling, including ectopic wing vein differentiation (Fig. 1F). Genetic downregulation of *Raf* and *ERK* suppresses Ras^G12V^-induced ectopic vein differentiation (data not shown). These findings indicate that *Ras1^G12V^*and *KRASB^G12V^* expressed from the endogenous *Ras1* locus are deleterious and drive excess signaling, even in the presence of a wild type allele. Genetic complementation assays reveal that all G12V mutant alleles fail to complement a *Ras1* deficiency (*Df(3R)by10*), and only *Ras1^G12V^* and *Flag-Ras1^G12V^*alleles partially complement a hypomorphic *Ras1* allele (*Ras1^e1B^*), with surviving adults displaying gain-of-function phenotypes (data not shown). In contrast, the *Drosophila* and human wild type *Ras* alleles complement with the two *Ras1* loss-of-function conditions, indicating that the reintegration resulted in functional copies of *Ras1*. The trans-heterozygous flies display normal phenotypes (Ras1^WT^) or *Ras1* loss-of-function phenotypes (Flag-Ras1^WT^, FRT Ras1, and Flag-KRASB^WT^), correlating with the amount of Ras protein levels produced from each allele. Notably, KRASB^WT^ rescued the *Ras1* knockdown phenotype (Fig. 1F), demonstrating that human KRASB can functionally substitute for its *Drosophila* homolog when expressed from the endogenous locus.

### Raf/MAPK pathway activation by *Ras^G12V^* forms

*Ras^G12V^* lines display dominant phenotypes characteristic of Ras/Raf/ERK pathway gain-of-function conditions. Pulldown assays using the Ras-Binding Domain (RBD) of Raf confirmed that Ras1^G12V^ and KRASB^G12V^ accumulate in the active, GTP-bound state, compared to the wild type protein (Fig. 2A-B). Despite this, overall levels of ERK phosphorylation were very similar in protein extracts from *Ras^G12V^* and normal *Ras* hemizygous and heterozygous larvae (Fig. 2D, E). We generated a Ras1^G12V^ variant harboring a mutation in the C-terminal CAAX motif (C186S, referred as Ras1^G12V-CAAX^), which prevents membrane anchoring (Michaelson *et al*, 2005). Ras1^G12V-CAAX^ is not accumulated in the active state (Fig. 2A, B), demonstrating that membrane localization is essential for nucleotide exchange of Ras1^G12V^ mutant proteins. Consistently, Ras1^G12V-CAAX^ did not induce Ras gain of function phenotypes (Fig. 2B) and was lethal in combination with *Ras* loss-of-function alleles.

**Figure 2.**
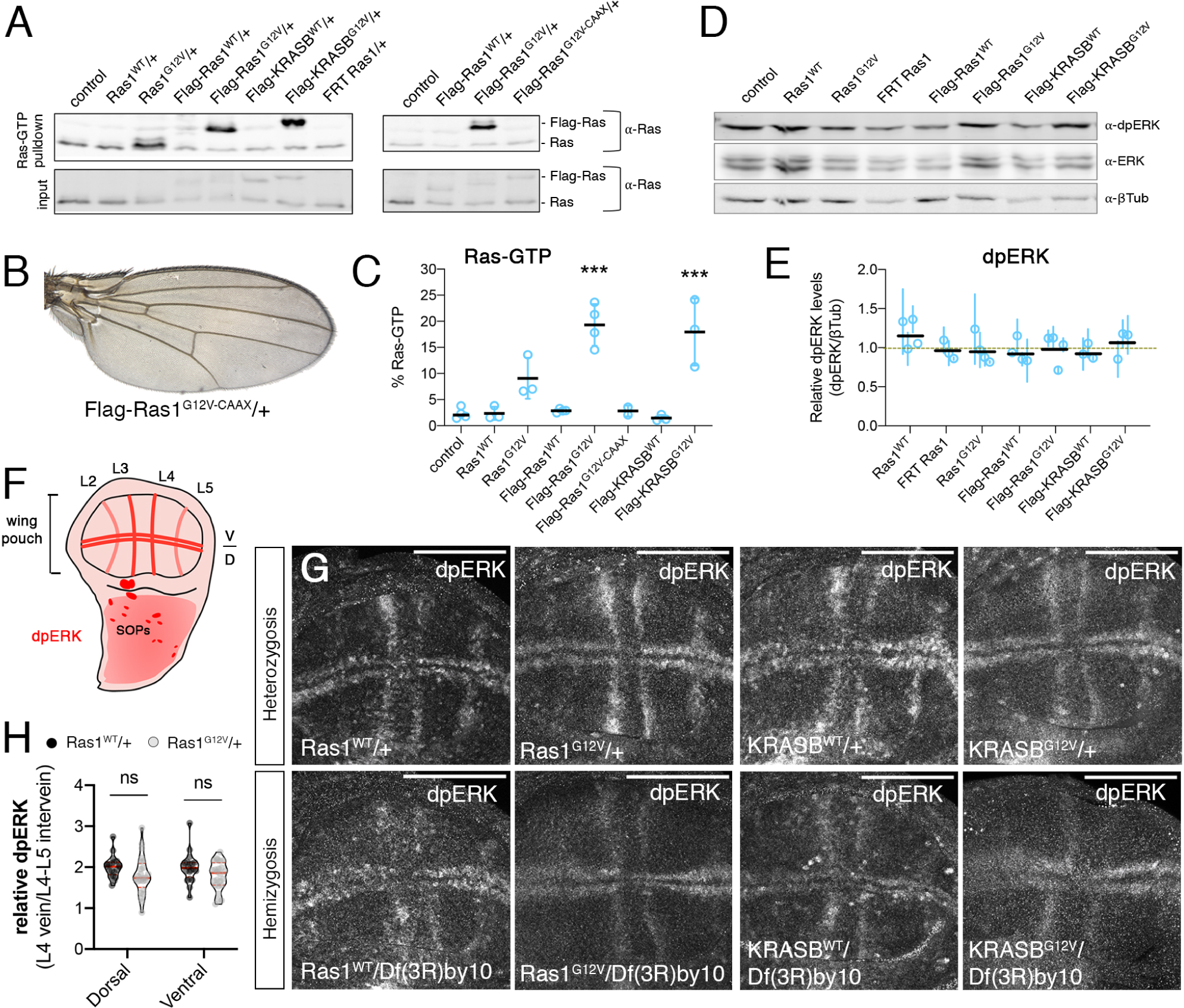
Analysis of Ras1^KO-Kin^ insertions. (A) Representative western blots of Raf-RAS binding domain (RBD) pulldowns detecting Ras-GTP from third instar larval extracts of the following genotypes: controls, *Ras1^WT^/+, Ras1^G12V^/+, Flag-Ras1^WT^/+, Flag-Ras1^G12V^/+, Flag-KRASB^WT^/+, Flag-KRASB^G12V^/+* and *FRT Ras1/+* (left) and control, *Flag-Ras1^WT^/+, Flag-Ras1^G12V^/+ and Flag-Ras1^G12V-CAAX^/+* (right). Each line shows the presence of Flag-Ras and Ras in the pulldown (above) and input (below). (B) Wing of Flag-Ras1^G12V-CAAX^/+ genotype showing normal wing size and pattern of veins. (C) Quantification of Ras-GTP (% Ras-GTP) corresponding to the western blots shown in A. The graph displays the mean ± SD of different biological replicates (dots). (D) Representative western blot from third instar larval extracts from hemizygous control (*Df(3R)by10/+*), *Ras1^WT^/Df(3R)by10, Ras1^G12V^/Df(3R)by10, Flag-Ras1^WT^/Df(3R)by10, Flag-Ras1^G12V^/Df(3R)by10, Flag-KRASB^WT^/Df(3R)by10* and *Flag-KRASB^G12V^/Df(3R)by10*. Blots were probed for dpERK, total ERK and βTub (loading control). (E) Quantification of dpERK levels relative to βTub corresponding to the western blots shown in D. No significant differences were observed between genotypes. Horizontal black lines indicate the mean of biological replicates. Each vertical blue line indicates the standard deviation (SD) of technical replicates, and the circles represents the mean of each biological replicate. (F) Schematic representation of a mature third instar larval wing disc indicating the pattern of dpERK accumulation by shades of red. In the wing pouch region, maximal accumulation of dpERK is detected in the developing wing veins (L2-L5), along two stripes abutting the dorso-ventral boundary (D/V) and in the precursor cells of the sensory organs (SOPs) in the dorsal notum region. (G) Immunostaining for dpERK in third instar larval wing imaginal discs of the following genotypes: *Ras1^WT^/+*, *Ras1^G12V^/+*, *KRASB^WT^/+* and *KRASB^G12V^/+* (top row from left to right) and *Ras1^WT^/Df(3R)by10*, *Ras1^G12V^/Df(3R)by10*, *KRASB^WT^/Df(3R)by10* and *KRASB^G12V^/Df(3R)by10* (bottom row, left to right). Scale bar: 100 μm. (H) Quantification of the ratio of dpERK levels between the vein L4 and the L4-L5 intervein territories in *Ras1^WT^/+* and *Ras1^G12V^/+* dorsal and ventral wing disc compartments. Violin plots display the median and the Q1 and Q3 interquartile ranges.

To examine Ras signaling *in situ*, we analyzed the distribution of dpERK in wing imaginal discs of *Ras1^G12V^* and *KRASB^G12V^* heterozygous and hemizygous larvae. In this epithelial tissue, ERK phosphorylation is normally detected at low levels in all cells, and displays a spatial pattern of maximal accumulation along the developing wing veins and in two stripes along the dorso-ventral margin of the wing pouch region (Fig. 2F-G) (Gabay *et al*, 1997). This pattern arises from the interplay between ligands and negative regulators of the EGFR pathway (Gabay *et al*, 1997), with low dpERK supporting growth and survival, while higher levels promoting vein differentiation (Diaz-Benjumea & Garcia-Bellido, 1990). Interestingly, the spatial pattern of dpERK maximal accumulation was not altered in the presence of Ras1^G12V^ or KRASB^G12V^, even when the activated mutant allele was the only form of Ras present in wing imaginal cells (*Ras1^G12V^/Df(3R)by10* and *KRASB^G12V^/Df(3R)by10*; Fig. 2G). Likewise, basal dpERK intensity in heterozygous mutant *Ras1^G12V^* wing discs was indistinguishable from controls (Fig. 2F). These results, which are fully consistent with the results of the genetic analyses and global dpERK accumulation, indicate that Ras^G12V^ expression cannot override the spatial control of ERK activation. Similarly, cell division and the incidence of cell death were not affected in *Ras1^G12V^*hemizygous wing discs (data not shown).

### Ras^G12V^ affects the temporal dynamics and basal levels of ERK phosphorylation

The acquisition of the spatial pattern of dpERK accumulation in the wing imaginal disc follows a precise temporal sequence, in which the first recognizable elements are two stripes along the dorso-ventral boundary, followed by activation in the presumptive L3, and finally L4 and L5 provein territories (Fig. 3A). In all cases examined, this temporal evolution occurs symmetrically between the dorsal and ventral wing compartments (Fig. 3A). However, when *Ras^G12V^*was expressed in the dorsal compartment using the conditional *FRT Ras1^WT^ FRT Ras1^G12V^* line (Fig. 3B), the accumulation of dpERK in its characteristic pattern occurs earlier in the dorsal (*Ras1^G12V^/+*; green) than in the ventral (*Ras1^WT^/+*; no GFP) compartment (Fig. 3C_1-3_). Despite of these different temporal dynamics, late third instar wing discs ultimately displayed a normal spatial pattern and levels of dpERK accumulation (Fig. 3C_3_, D). A similar acceleration of pathway activation was observed for human KRASB^G12V^ (Fig. 3E-G) and also when mutant proteins were restricted to the posterior compartment of the wing discs (data not shown). These results indicate that Ras^G12V^ promotes precocious activation of the Ras/Raf/ERK pathway within its normal spatial domains.

**Figure 3.**
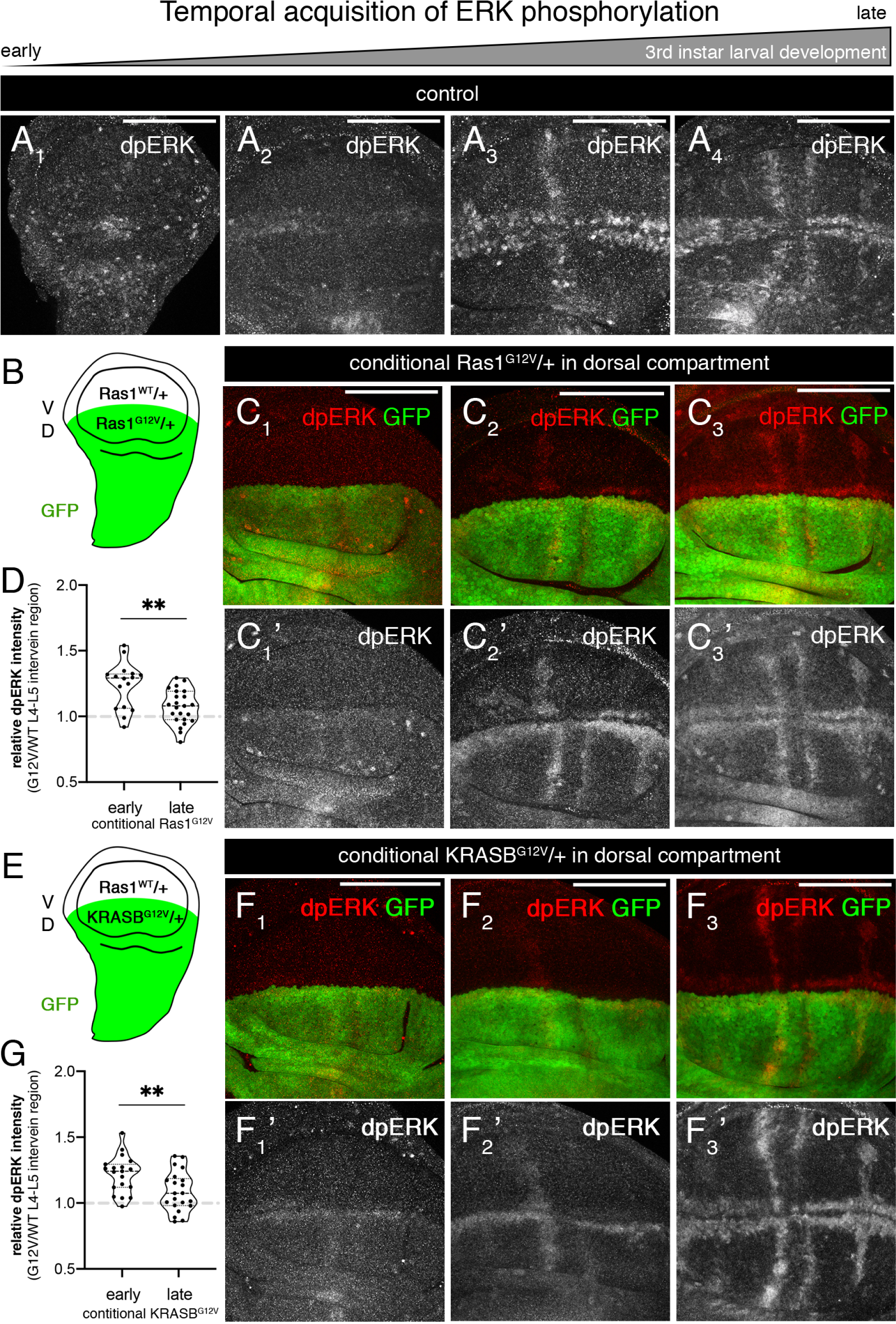
Temporal evolution of dpERK accumulation in the wing disc. (A) Accumulation of dpERK in third instar wing imaginal discs, from younger (A_1_) to older (A_4_). (B) Schematic representation of a mature wing disc of *ap-Gal4 UAS-GFP/UAS-FLP; FRT Ras1^WT^ FRT Ras1^G12V^/+* genotype showing the generation of dorsal *Ras1^G12V^/+* (GFP positive) and ventral *Ras1^WT^/+* (GFP negative) territories. (C) Expression of GFP (green in C_1_-C_3_) and dpERK (red in C_1_-C_3_ and white in C_1_’-C_3_’) in progressively older third instar wing discs of *ap-Gal4 UAS-GFP/UAS-FLP; FRT Ras1^WT^ FRT Ras1^G12V^/+* genotype. (D) Quantification of dpERK levels in dorsal (Ras1^G12V^/+) relative to ventral (Ras1^WT^/+) compartments in the L4-L5 intervein region of early and late third instar wing disc. Violin plots display the median and the Q1 and Q3 interquartile ranges. (E) Schematic representation of a mature wing disc of *ap-Gal4 UAS-GFP/UAS-FLP; FRT HA-Ras1^WT^ FRT Flag-KRASB^G12V^/+* genotype showing dorsal KRASB^G12V^/+ (GFP positive) and ventral Ras1^WT^/+ (GFP negative) territories. (F) Expression of GFP (green in F_1_-F_3_) and dpERK (red in F_1_-F_3_ and white in F_1_’-F_3_’) in progressively older third instar wing discs of *ap-Gal4 UAS-GFP/UAS-FLP; FRT HA-Ras1^WT^ FRT Flag-KRASB^G12V^/+* genotype. (G) Quantification of dpERK levels in dorsal (KRASB^G12V^/+) relative to ventral (Ras1^WT^/+) compartments in the L4-L5 intervein region of early and late third instar wing disc. Violin plots display the median and the Q1 and Q3 interquartile ranges. Scale bar: 100 μm.

We also analyzed homozygous *Ras1^G12V^* cells generated in the imaginal wing epithelium (Fig. 4A). These cells reached maximal dpERK levels more efficiently than adjacent heterozygotic *Ras1^G12V/+^*or wild type *Ras1^+/+^* cells (Fig. 4B, D). Interestingly, *Ras1^G12V/G12V^* clones were twice the size of wild type twins (Fig. 4B-C) and homozygotic *Ras1^G12V^*clones presented a 30% increase in the basal levels of dpERK in intervein regions (Fig. 4B, D). These results suggest that Ras1^G12V^ increases basal ERK phosphorylation in regions of low activity, thereby conferring a growth advantage, while maximal signaling remains constrained by upstream regulatory inputs from EGFR. Consistently, we do not observe ectopic expression of the vein territory marker Delta at intervein regions in *Ras1^G12V^*homozygous clones (Fig. 4E), and adult wings do not display the formation of ectopic vein tissue.

**Figure 4.**
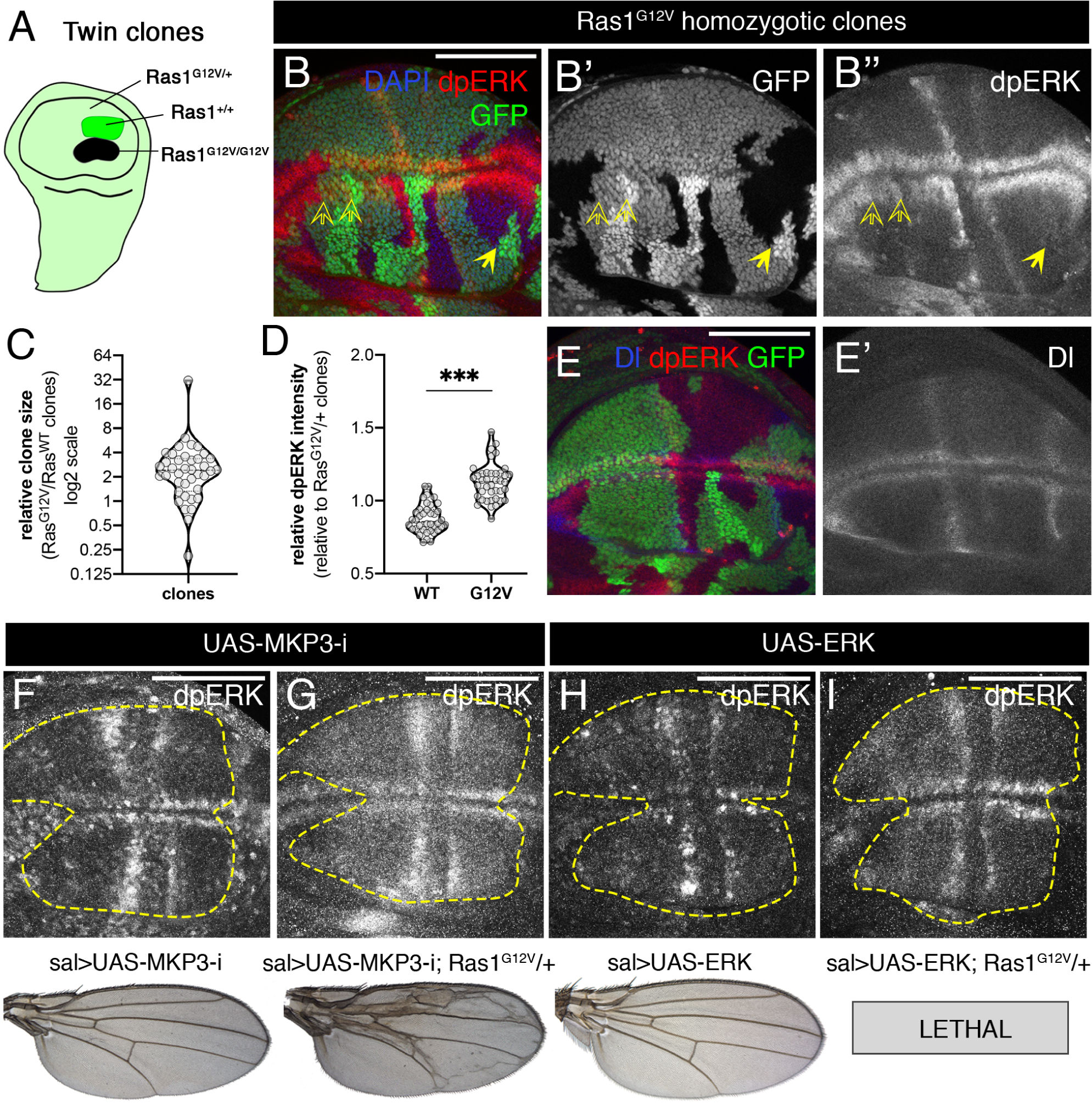
Consequences of Ras1^G12V^ gene dosage and interactions with MKP3 and ERK levels. (A) Schematic representation of a mature wing disc of *hs-FLP/+; FRT82B Ras1^G12V^/FRT82B Ubi-GFP* genotype showing the generation of wild type *Ras1* (Ras1^+/+^; bright GFP) and *Ras1^G12V^*(Ras1^G12V/G12V^; GFP negative) homozygous clones in a background of Ras1^G12V/+^ heterozygous cells (light green). (B) Representative wing disc bearing Ras1^+/+^ and Ras1^G12V/G12V^ clones showing anti-dpERK staining (red), GFP expression (green) and DAPI staining (blue). Single-channel images for GFP and dpERK are shown in B’ and B’’ respectively. (C) Quantification of the area occupied by *Ras1^G12V/G12V^* clones relative to the area occupied by *Ras1^+/+^* twin clones. The violin plot displays the median and the Q1 and Q3 interquartile ranges. (D) Quantification of relative dpERK intensity in *Ras1^G12V/G12V^* and *Ras1^+/+^* twin clones relative to heterozygotic *Ras1^G12V/+^* cells in intervein regions. Violin plots display the median and the Q1 and Q3 interquartile ranges. (E-E’) Expression of Dl (red in E, white in E’) in a wing disc of *hs-FLP/+; FRT82B Ras1^G12V^/FRT82B Ubi-GFP* genotype with homozygous wild type *Ras1* (Ras1^+/+^; bright GFP) and *Ras1^G12V^* (Ras1^G12V/G12V^; GFP negative) homozygous clones in a background of *Ras1^G12V/+^* heterozygous cells (light green). (F-G) Genetic interaction between endogenous *Ras1^G12V^* and *MKP3*. The adult wings of *sal^EPv^-Gal4 UAS-GFP/UAS-MKP3-RNAi* (sal>UAS-MKP3-i) are normal (F), whereas the wings of *sal^EPv^-Gal4 UAS-GFP/UAS-MKP3-RNAi; Ras1^G12V^/+* (sal>UAS-MKP3-i; Ras1^G12V^/+) display excess of vein material (G). The corresponding third instar wing imaginal discs showing the accumulation of dpERK are shown above each wing, and the area of *sal^EPv^-Gal4* expression is circled by yellow dotted lines. (H-I) Genetic interaction between endogenous *Ras1^G12V^* and increased ERK levels. The adult wings of *sal^EPv^-Gal4 UAS-GFP/UAS-ERK* (sal>UAS-ERK) are normal (F), whereas the combination *sal^EPv^-Gal4 UAS-GFP/UAS-ERK; Ras1^G12V^/+* (sal>UAS-ERK; Ras1^G12V^/+) is pupal lethal. The corresponding third instar wing imaginal discs showing the accumulation of dpERK are shown above, and the *sal^EPv^-Gal4* domain is circled by yellow dotted lines. Scale bar: 100 μm.

To assess whether Ras^G12V^ enhances basal ERK phosphorylation under sensitized conditions, we examined genetic backgrounds prone to detect weak alterations in ERK activation. Knockdown of the ERK dual-specificity phosphatase *MKP3* (Fig. 4F) or ectopic expression of ERK (Fig. 4H) did not affect wing vein development (Fig. 4F, H). Consistently, the localization of dpERK at maximal levels was restricted to the proveins in the corresponding wing imaginal discs (Fig. 4F and H). By contrast, *MKP3* knockdown or *ERK* overexpression in a *Ras1^G12V^* background increases the background levels of ERK phosphorylation without altering the localization of maximal dpERK in the developing veins (Fig. 4G and I). These increases induced ectopic vein formation in the proximity of the normal veins (*UAS-MKP3-RNAi; Ras1^G12V^/+*; Fig. 4G) and pupal lethality (*UAS-ERK; Ras1^G12V^/+*; Fig. 4I). Taken together, these results suggest that *Ras1^G12V^* does not disrupt the normal spatial regulation of Ras/Raf/ERK pathway activation in the wing disc. Instead, *Ras1^G12V^* advances the timing of maximal pathway activation and elevates basal ERK phosphorylation.

### Modulation of Ras1^G12V^ signaling by upstream mediators

The normal spatial pattern of dpERK observed in Ras^G12V^ suggests that its activity is controlled by upstream regulators, including GAP and GEF proteins and the EGFR. We found that *Ras1^G12V^* suppress the vein loss caused by excess of GAP (Fig. 5A-B) and knockdown of GEF (Fig. 5C-D). Consistently, the accumulation of dpERK in the developing veins was rescued in the corresponding wing imaginal discs (Fig. 5A’-D’). Thus, Ras1^G12V^ is independent of GAP, as expected, but can compensate for a reduction in SOS. In addition, *Ras1^G12V^* combined with overexpression of EGFR caused a synergistic differentiation of ectopic veins (Fig. 5E-F), suggesting that endogenous *Ras1^G12V^*can respond to EGFR activation *in vivo*. Finally, *Ras1^G12V^*rescued the size of wings with reduced expression of EGFR (Fig. 5G-H), but not the loss of veins or the accumulation of dpERK (Fig. 5G-H, G’-H’).

**Figure 5.**
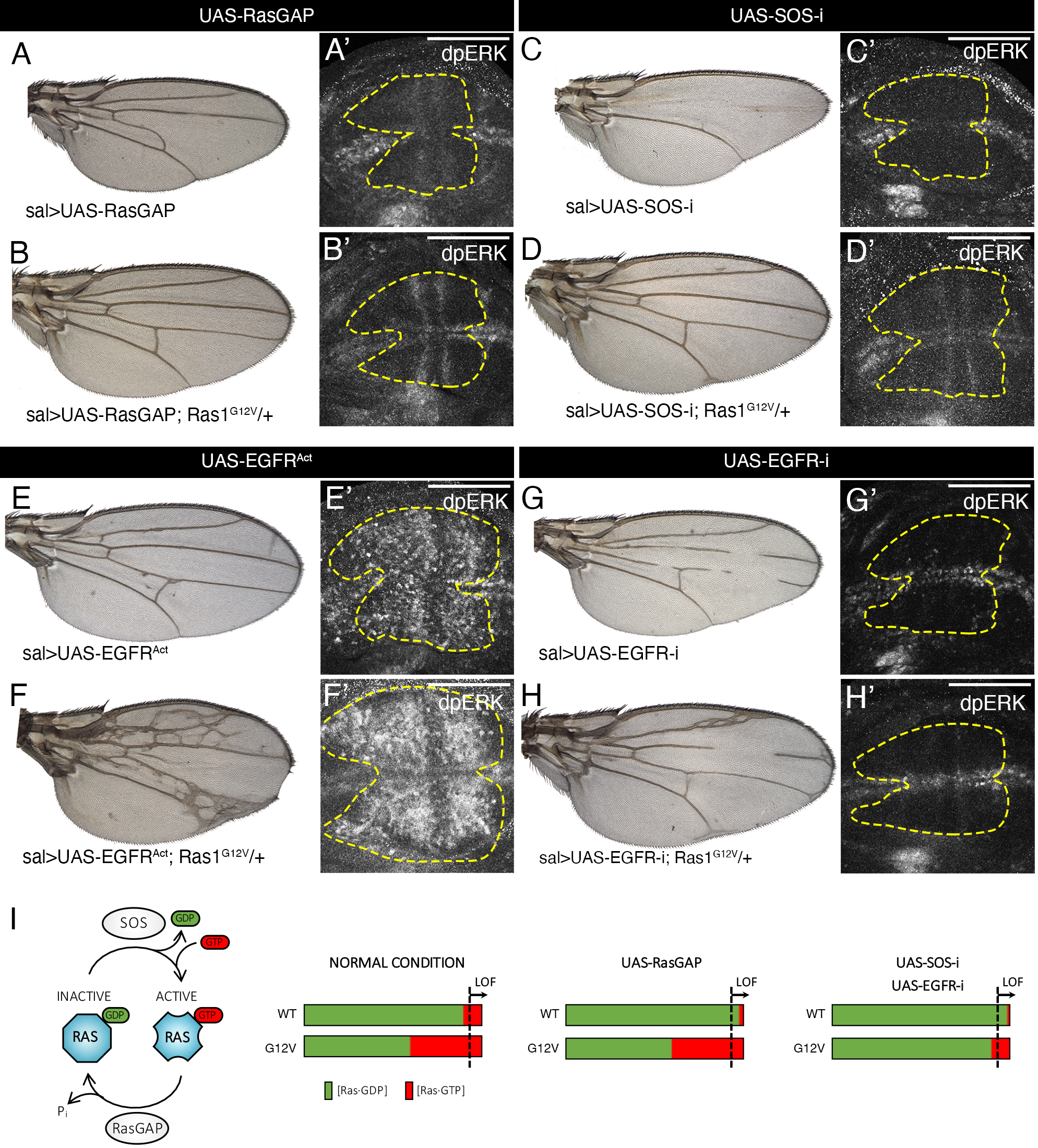
Interactions of Ras1^G12V^ with its direct upstream regulators RasGAP, SOS and EGFR. (A-A’) The overexpression of GAP (*sal^EPv^-Gal4 UAS-GFP/UAS-RasGAP*) results in a loss of L4 vein phenotype (A) and reduced dpERK accumulation in the corresponding third instar wing imaginal disc (A’). (B-B’) The heterozygosity of *Ras1^G12V^* (*sal^EPv^-Gal4 UAS-GFP/UAS-RasGAP; Ras1^G12V^/+*) rescues the phenotype of *RasGAP* overexpression (B), and the reduction of dpERK expression (B’). (C-C’) The knockdown of *SOS* (*sal^EPv^-Gal4 UAS-GFP/UAS-SOS-RNAi*) results in a loss L2-L4 veins C) and reduced dpERK accumulation in the corresponding third instar wing imaginal disc (C’). (D-D’) The heterozygosity of *Ras1^G12V^* (*sal^EPv^-Gal4 UAS-GFP/ UAS-SOS-RNAi; Ras1^G12V^/+*) rescues the loss of function phenotype caused by knockdown of *SOS* (D), and the reduction of dpERK accumulation (D’). (E-E’) Ectopic expression of activated EGFR (*sal^EPv^-Gal4 UAS-GFP/+; UAS-EGFR^λAtop^/+*) causes a weak excess of vein tissue (E) and a generalized phosphorylation of ERK (E’). (F-F’) Excess of activated EGFR in a *Ras1^G12V^* background (*sal^EPv^-Gal4 UAS-GFP/+; EGFR^λAtop^/Ras1^G12V^*) causes a strong excess of vein tissue (F), and increases the accumulation of dpERK (F’). (G-G’) Knockdown of EGFR (*sal^EPv^-Gal4 UAS-GFP/+; UAS-EGFR-RNAi/+*) causes a weak loss of L3 and L4 vein tissue (G) and loss of ERK phosphorylation (G’). (H-H’) Knockdown of EGFR in a *Ras1^G12V^*background (*sal^EPv^-Gal4 UAS-GFP/+; UAS-EGFR-RNAi/Ras1^G12V^*) retain the loss of vein phenotype (H), and the loss of dpERK (H’). (I) Schematic representation of Ras activation in genetic backgrounds with altered expression of *SOS*, *RasGAP* and *EGFR* in *Ras1* wild type (WT) and *Ras1^G12V^* heterozygous backgrounds (G12V). Green and red shading represent estimated fractions of Ras bound to GDP (Ras-GDP) or GTP (Ras-GTP) in normal conditions (left), upon over expression of *RasGAP* (UAS-RasGAP; center) and upon knockdown of *SOS* (UAS-SOS-RNAi) or EGFR (UAS-EGFR-RNAi). Scale bar: 100 μm.

To directly evaluate the requirement of EGFR activity for Ras^G12V^ signaling, we analyzed cells with no *EGFR* function. In these experiments, we compared the behavior of *EGFR* mutant cells growing in wild type *Ras1* or in *Ras1^G12V^* genetic backgrounds (Fig. 6A). We used the *EGFR* alleles *EGFR^f24^*, *EGFR^f3^*, *EGFR^f2^*and *EGFR^f5^*. All these alleles fail to complement with each other, and are described as strong *EGFR* loss of function conditions (Price *et al*, 1989; Clifford & Schupbach, 1994). In all cases we found that *EGFR* mutant clones exhibited reduced growth (Fig. 6B-C, E-F and data not shown) and failed to reach high levels of ERK phosphorylation in the DV border and proveins (Fig. 6C’’ and data not shown). Strikingly, *EGFR* mutant clones were readily recovered when growing in a *Ras1^G12V^*heterozygous background (Fig. 6 B, D, E, G and data not shown). Clone size was fully rescued in *EGFR^f24^* (Fig. 6B), *EGFR^f2^* and *EGFR^f5^*(not shown) and partially rescued (70%) for the *EGFR^f3^* allele (Fig. 6E). Interestingly, high levels of ERK phosphorylation were still lost in all *EGFR* mutant clones growing in a *Ras1^G12V^* heterozygous background (Fig. 6D. G and data not shown), suggesting that EGFR activity is essential for maximal levels of dpERK in the presence of *Ras1^G12V^*. In the corresponding wings, we noticed a combination of loss of normal vein stretches associated to the formation of ectopic vein tissue close to the position normally occupied by the vein (Fig. 6H), as described previously (Diaz-Benjumea & Garcia-Bellido, 1990). Collectively, these observations suggest that Ras^G12V^ can sustain low-level Raf/ERK pathway activation sufficient for growth of wing disc epithelial cells independently of *EGFR.* However, full pathway activation, needed to promote vein cell differentiation, remains EGFR dependent. (Fig. 6I).

**Figure 6.**
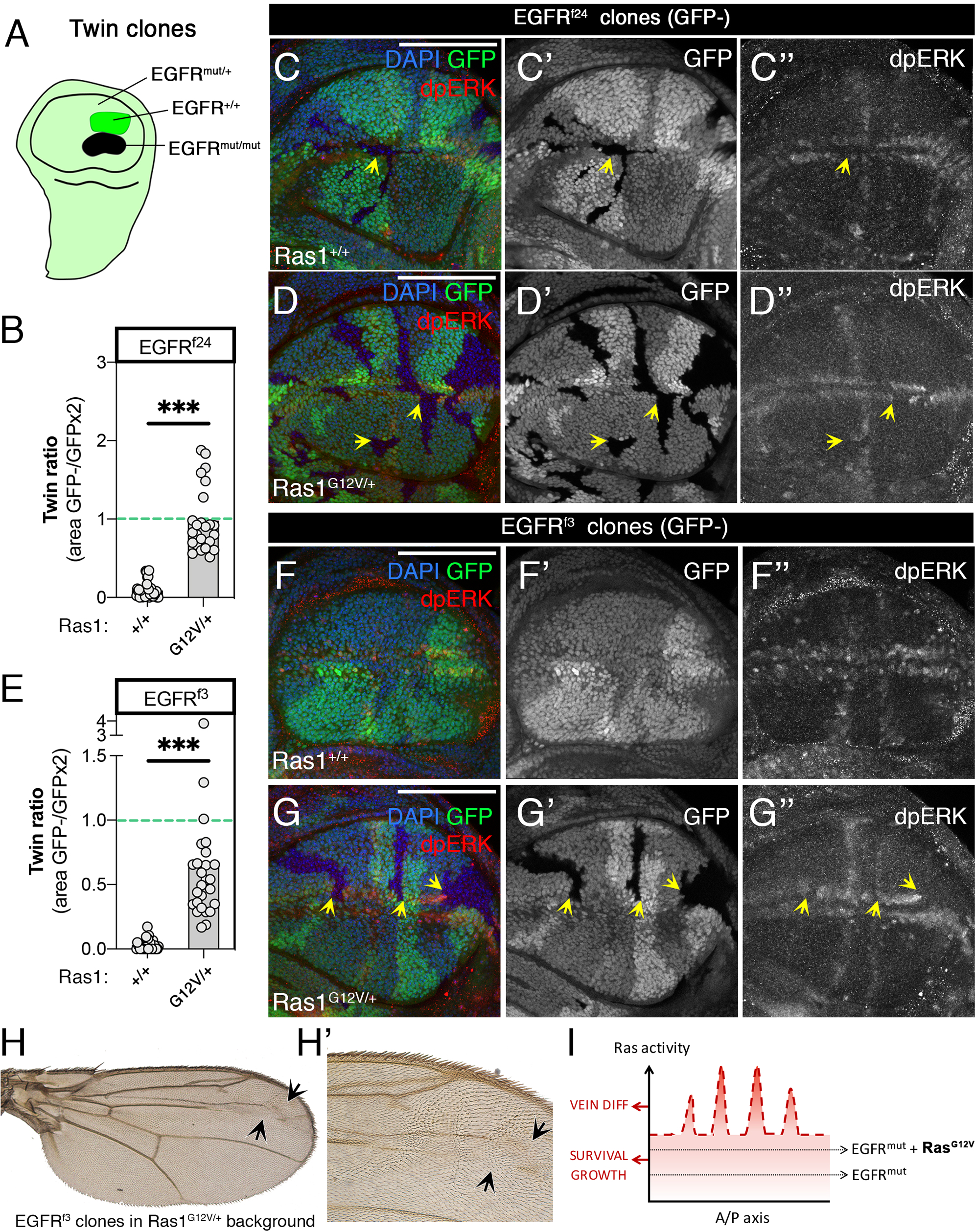
Clonal analysis of *EGFR* mutant alleles in a *Ras1^G12V^* background. (A) Schematic representation of a late third instar disc of *hs-FLP; FRT42D Ubi-GFP/FRT42D EGFR^mut^* genotype allowing the generation of twin clones of *FRT42D EGFR^mut^* (GFP^-/-^) and *FRT42D Ubi-GFP* (GFP^+/+^) genotypes. Background heterozygous cells have only one copy of *Ubi-GFP*, and are represented in pale green (GFP^+/-^). (B) Quantification of the area ratio between *EGFR^f24^*mutant (GFP^-/-^) and wild type (GFP^+/+^) twin clones growing in a wild type *Ras1* background (+/+; left) or in a *Ras1^G12V^/+* background (G12V/+). Bar graphs display the mean value. (C-C’’) Representative example of mutant *EGFR^f24^* and wild type *EGFR* twin clones generated in a *Ras1* wild type background. The expression of GFP is in green in C and grey in C’. The accumulation of dpERK is in red in C and grey in C’’. (D-D’’) Representative example of mutant *EGFR^f24^* and wild type *EGFR* twin clones generated in a *Ras1^G12V^/+* background. The expression of GFP is in green in D and grey in D’. The accumulation of dpERK is in red in D and in grey in D’’. (E) Quantification of the area ratio of *EGFR^f3^* mutant (GFP^-/-^) and wild type (GFP^+/+^) twin clones growing in a wild type *Ras1* background (+/+; left) or in a *Ras1^G12V^/+* background (G12V/+). Bar graphs display the mean value. (F-F’’) Representative example of mutant *EGFR^f3^* and wild type *EGFR* twin clones generated in a *Ras1* wild type background. The expression of GFP is in green in F and grey in F’. The accumulation of dpERK is in red in F and in grey in F’’. (G-G’’) Representative example of mutant *EGFR^f3^* and wild type *EGFR* clones generated in a *Ras1^G12V^/+* background. The expression of GFP is in green in G and grey in G’. The accumulation of dpERK is in red in G and in grey in G’’. Note the loss of high levels of dpERK in *EGFR^f24^*(C’’-D’’) and *EGFR^f3^* (G’’) homozygous cells (yellow arrowheads). The staining of DAPI in shown in blue in C-G. (H) Adult wing of *hs-FLP; FRT42D Ubi-GFP/FRT42D EGFR^f3^; Ras1^G12V^/+* showing loss of L3 vein (arrowheads). H’ is a higher magnification of the distal part of the wing shown in H. (I) Schematic representation of dpERK levels (red line) along the A/P axes of the wing disc indicating two different cellular responses: vein differentiation and survival/growth. Effects of loss of *EGFR* function in *Ras1* wild type (EGFR^mut^) and *Ras1^G12V^* (EGFR^mut^ + Ras1^G12V^) backgrounds. Scale bars: 100 μm.

## Discussion

The levels of Ras activity are determined by a variety of functional parameters, including expression levels of specific Ras isoforms and their mutational states (Muñoz-Maldonado *et al*, 2019). In addition, the translation of a particular quantity of Ras activation into a biological outcome relies on the cellular type, the cell developmental state and the occurrence of additional genetic or physiological alterations. In the framework of tumoral growth, these notions have been encapsulated in the “sweet-spot” model, which proposes that for each cellular and genetic setting there is an optimal level of Ras activity that promotes tumor formation (Li *et al*, 2018). The relevance of Ras activity levels imposes that animal or cellular models used in Ras research must recapitulate as closely as possible the conditions found in pathological conditions, thereby enabling pharmacological, genetical or biochemical research of maximal therapeutic relevance.

In this work we developed an experimental platform allowing the evaluation of different mutant versions of Ras and different Ras isoforms expressed from the *Ras1* regulatory sequences, and we characterized how Ras1^G12V^ and KRASB^G12V^ signal *in vivo*. A major advantage of *Drosophila* is that it contains only one *Ras* gene, and no redundancy in downstream components of the pathway. Furthermore, EGFR is the only tyrosine kinase receptor that controls Ras activity in the wing disc, and consequently the interpretation of the results does not require to call for complex feedback loops affecting other Ras isoforms or tyrosine kinase receptors.

The evaluation of wild type and G12V mutant Ras1 and KRASB proteins indicates that KRASB can functionally replace Ras1 *in vivo*, and that both proteins exhibit comparable pathway activation and genetic behavior in complementation tests. They also produce a similar spectrum of phenotypes, allowing the extrapolation of results obtained in *Drosophila* Ras1^G12V^ to human KRASB^G12V^. Despite targeting the endogenous *Ras1* locus, we only obtained Ras^G12V^ lines presenting lower-than-normal Ras protein levels. Embryonic expression of the oncogenic forms correlates with compromised viability, which together with the impossibility of obtaining transgenic strains with normal Ras^G12V^ protein levels indicates dominant lethality of this allele, consistent with findings in other experimental models. For example, murine, models expressing oncogenic *Kras* or *Nras* mutations during embryonic development result in dominant lethality (Guerra *et al*, 2003; Tuveson *et al*, 2004; Wang *et al*, 2011), which is overcome when the protein levels of mutant Nras are reduce up to 25% (Wang *et al*, 2011). Taken together, our results indicate that *Ras1^G12V^* behaves as a dominant lethal allele when expressed at normal levels, and as a dominant allele with phenotypes affecting larval and adult viability when expressed at reduced levels.

### Relationships of endogenous Ras^G12V^ and ERK activation

The Ras^G12V^ protein expressed at near physiological levels promotes phenotypes consistent with Raf/MAPK signaling pathway overactivation, without modifying overall levels of ERK phosphorylation. Similarly, mice models with endogenous ras activating mutations display developmental defects and tumorigenic behaviors, but Raf/ERK pathway activation is not overly altered in tissues and cell cultures derived from them (Guerra *et al*, 2003; Braun *et al*, 2003; Schuhmacher *et al*, 2008; Chen *et al*, 2009). More recently, the use of FRET-based biosensors with enhanced sensitivity to define ERK activation dynamics suggests that activated Ras isoforms raise baseline ERK activity but reduce peak responses to EGF (Gillies *et al*, 2020; Ponsioen *et al*, 2021). These observations suggest that the phenotypic effects promoted by Ras^G12V^ are more likely related to perturbation at baseline levels of downstream pathway activation and/or inappropriate activation/inactivation dynamics. Consistently, and in contrast to ectopically expressed *Ras1^G12V^* and *KRASB^G12V^*, endogenous Ras1^G12V^ or KRASB^G12V^ only cause a weak increase in the basal levels of ERK phosphorylation *in situ,* with peaks of pathway activation present at the normal locations that are acquired earlier by mutant cells compared to wild type cells. Thus, Ras^G12V^ mutant cells display increased sensitivity to EGFR stimulation to promote ERK activation, as manifested in mosaic experiments and combinations in sensitized genetic backgrounds. These results indicate that ERK activity is determined by at least two regulatory layers: the spatial distribution of the upstream EGFR regulators, such as ligands expression, and the level and mutational status of Ras. Overall, Ras^G12V^ mutants cause only moderate increases in ERK phosphorylation, likely constrained by the normal regulatory mechanisms that preserve the spatial and temporal fidelity of pathway activation.

### Ras^G12V^ is not a constitutive active mutant and requires EGFR activity

Ras activity is controlled in the wing epithelium by EGFR signaling, which is regulated by the spatial and temporal expression of different EGFR ligands (Gabay *et al*, 1997), and fine-tuned by positive and negative feedback loops (Oda *et al*, 2005; Freeman, 2000). The conservation of the normal pattern of ERK activation observed in heterozygotic, hemizygotic, and homozygotic conditions indicates that Ras1^G12V^ and KRASB^G12V^ activation still relies on upstream stimulation, regardless of protein levels or the presence of wild type *Ras1*. Consistently, Ras1^G12V^ requires membrane anchorage to increase the pool of Ras-GTP molecules, and responds more efficient than wild type Ras to EGFR stimulation. Importantly, Ras1^G12V^ can sustain basal ERK activity and promote the survival of EGFR mutant cells, thus being independent from EGFR activity to sustain certain cellular responses associated with low-levels of pathway activation. At this point we cannot rule out whether other RTK, such as for example the Insulin receptor, which is expressed in all wing imaginal disc cells, can activate Ras^G12V^ in *EGFR* mutant cells to provide basal activation of the Raf/ERK pathway through mutant *Ras1*. In any case, maximal levels of the pathway are strictly dependent on EGFR activity and in this context the G12V mutation cannot be considered as constitutively activated. Whether this dependence is shared by other activating *Ras* mutations will require specific evaluation.

There is a wealth of information concerning the dependence of activating Ras mutations on EGFR activity. On the one hand, KRAS mutations associated with poor prognosis in colorectal cancers (CRC) and non–small-cell lung cancer are strong negative predictors of response to anti-EGFR therapies (Thein *et al*, 2021), and *KRAS* mutations are associated with acquired resistance to anti-EGFR treatment in CRC and organoids harboring Ras activating mutations (Verissimo *et al*, 2016). These findings suggest that Ras activation can occur independently of upstream signals. In contrast, EGFR RNAi is highly effective to target *KRAS^G12V^*in different tumoral models (Stanland *et al*, 2025), and KRAS^G12C^ inhibitors synergize with EGFR inhibitors in tumoral cell lines (Ryan *et al*, 2019). Furthermore, the elimination of EGFR in genetic mouse models for lung and pancreatic cancer suppress the growth of *Kras^G12V^* and *Kras^G12D^* mutant tumors (Navas *et al*, 2012; Moll *et al*, 2018; Ardito *et al*, 2012; Blasco *et al*, 2019) and amplification of other RTKs is a resistance mechanism in *KRAS^G12C^*-mutant tumors after KRAS^G12C^ inhibitor treatment (Akhave *et al*, 2021). These results point out to a key role of the EGFR receptor to activate ERK signaling in the presence of activated Ras mutations, which might involve the activation of both wild type and mutated Ras proteins. Consequently, a combinatorial therapy against KRAS^G12C^ (adagrasib) and EGFR (cetuximab) was recently approved by FDA for treating CRC (FDA, 2024). We hypothesize that the dependence of Ras^G12V^ mutants on EGFR activity relies on the cellular context, including the type of tyrosine kinase receptors present, the level at which Ras activity promote specific cellular responses and the role of the remaining wild type Ras isoforms, which through complex feedback interactions can impact in the final outcome of the pathway. In this manner, the efficacy of EGFR inhibition in the presence of Ras^G12V^ activating mutation may depend on the availability of other TKR receptors able to activate Ras^G12V^ and in the level of EGFR inhibition, where partial loss may lead to enough Ras signaling beyond a threshold able to support tumor development and progression depending on the cellular context.

## Methods

### *Drosophila* stocks and genetics

We used the *Gal4* lines *sal^EPv^-Gal4* (wing region located between the vein L2 and the intervein L4/L5; Cruz et al., 2009)*, nos-Gal4* (BL4442), *en-Gal4* and *hh-Gal4* (posterior compartment of the wing disc) and *ap-Gal4* (dorsal compartment of the wing disc). We also used the following UAS lines: *UAS-MKP3-RNAi* (VDRC23911/GD), *UAS-ERK-HA* (Molnar and de Celis 2013), *UAS-ERK-RNAi* (VDRC35641/GD), *UAS-Raf-RNAi* (VDRC20909/GD), *UAS-GFP* (Ito *et al*., 1997), *UAS-FLP*, *UAS-Ras1-RNAi* (VDRC106642/KK), *UAS-Ras1^V12^* (BL64196), *UAS-Flag-Ras1^V12^*(Vega-Cuesta et al. 2020), *UAS-RasGAP* (*EP160*; Molnar et al. 2006), *UAS-Sos-RNAi* (BL34833), *UAS-EGFR^WT^* (BL5268), *UAS-EGFR^α−Top^* (Queenan et al. 1997), *UAS-EGFR-RNAi* and *UAS-EGFR^DN^*(BL5634). We also used a genetic deficiency for the *Ras1* locus (*Df(3R)by10*; BL1931), the *Ras1* alleles *Ras1^e1B^* (BL5689), *Ras1^λ1c40b^* (Hou et al. 1995), the *EGFR* alleles *EGFR^f24^* (BL51296), *EGFR^f2^* (kindly provided by Trudi Shupard and Sonsoles Campuzano), *EGFR^f3^* (BL34043), and *EGFR^f5^*(BL34504), and the *Sos* allele *Sos^k05224^* (BL10566). The sequence of *EGFR* mutant alleles revealed that the expected mutation on *EGFR^f2^*was not present in any of the *EGFR^f2^* lines we had available. The *EGFR^f2^*, *EGFR^f3^*, and *EGFR^f5^*alleles were recombined on a *FRT42D* chromosome. Twin spot clones were generated using the lines *hs-FLP; Ubi-GFP, FRT40A/ CyO* and *hs-FLP; FRT42D Ubi-GFP/CyO*. Clone induction was performed at 72 h AEL by heat shock at 37 °C for 1 h. *Ras1^KO^* and *Ras1^KO-Kin^* flies were generated using the strains *nanos-ϕC31*, *hs-Cre/CyO; TM2/TM6b*, *vasa-Cas9* (BL55821) and *hs-FLP; If/CyO; MKRS/TM6b*. Flies were raised at 25 °C unless otherwise stated.

### Generation of *Ras1^KO^* and *Ras1^Kin^* strains

We generate *Ras1 knockout* (*Ras1^KO^*) transgenic lines by CRISPR-Cas9-enhanced homologous recombination to replace the *Ras1* coding sequence with an *attP* landing site (Baena-Lopez *et al*, 2013). *Ras1^Kin^* lines were generated from the *Ras1^KO^* line using *ϕC31* mediated recombination. Briefly, *Drosophila Ras1* genomic DNA from *vasa-Cas9* flies, *Ras1* cDNA (RE53955, DGCR) and human *KRASB* cDNA (SC109374, OriGene Technologies) were cloned into reintegration vectors (*RIV^white^*and *RIV^FRTMCSFRTMCS3^*) (Baena-Lopez *et al*, 2013) in either wild-type or G12V mutant forms and tagged or not with the Flag or HA epitopes at their 5’ end. All *Ras1 knock-in* constructs included the 5′ and 3′ UTRs deleted in the *Ras1^KO^* line (Fig. 1A). Fig. 1A summarizes all *Drosophila Ras1* and human *KRASB* knock-in lines generated. The reintegration vectors were injected in embryos of *nanos-phiC31; Ras1^KO^/+* genotype. All the *knock-out* and *knock-in* lines generated were confirm by DNA sequencing.

### Western Blots

In brief western blotting was performed from larval lysates (10 larvae per replicate condition) washed in cold PBS (4 °C), dried, and stored at −80 °C until use. For each sample, 75 µg of total protein were resolved by SDS-PAGE using 7.5% gels (phosphoprotein detection) or 10% gels (Ras detection). All images were processed with Image Fiji and Adobe Photoshop CC 2017. Western blots were repeated at least for three biological replicates.

### GST-Raf.RBD pull-downs assays

Pulldown assays were carried out following the protocols described in ref (Baker & Rubio, 2021). Protein samples were loaded in 10% SDS-PAGE. Between 3 and 5 biological replicates were performed, except for *Flag-Ras1^G12V-CAAX^*genotype (2x).

### Immunofluorescence in larval tissues

Imaginal wing discs were dissected, fixed, and stained as described in ref (de Celis, 1997). Confocal images were taken in a LSM710 confocal microscope (Zeiss). All images were processed with ImageJ 2.14.0 (NIH, USA) and Adobe Photoshop CC 2017.

### Pupal lethality and viability assays

Pupal lethality was determined as the percentage of dead pupae in the progeny of 10-20 control virgin females (*nanos-ϕC31*) and 5-12 males of the *Ras1^KO-Kin^/TM6b* genotype. Viability was assessed by comparing the number of pupae and adults of *Ras1^KO-Kin^/+* to those of *TM6b/+* siblings within each vial. We analyzed a minimum of 100 individuals per genotype in each experiment.

### Survival assays

20 newly emerged flies were transferred to fresh vials and subsequently moved to new vials three times per week. Survival was monitored every three days by recording the number of living individuals. At least 100 flies were analyzed per genotype, except for control males (n=79), *Flag-KRASB^G12V^/+* males (n=35), and *Flag-Ras1^WT^/+* females (n=89).

### Quantifications and statistical analysis

All numerical data were collected and processed in Microsoft Excel (Microsoft Inc.). Measurements of mitotic index were done with Adobe Photoshop CC 2017. Fluorescence or band intensity in immunostainings or western blots, respectively, and percentage of area positive for Dcp1 staining were calculated in ImageJ2. Statistical analyses were done with GraphPad Prism 8. Prior to the statistical analysis we performed normality tests (Shapiro-Wilk). Appropriate analysis for further comparisons were performed (Unpaired t-test, Welch’s t test, Mann-Whitney, Ordinary one-way ANOVA or Multiple comparison) using GraphPad Prism 8. The statistical analyses of survival curves were carried out using Gehan-Breslow-Wilcoxon test. We consider that there is a significant difference when the p-value is lower than 0.001 (***), 0.01 (**) or 0.05 (*). Graphic representations were done using GraphPad Prism 8. Box plots show the median and the Q1 and Q3 quartiles. Column bar graphs show mean with standard deviation (SD). Violin plots show the median and the Q1 and Q3 quartiles.

## Data Availability

All data generated or analyzed during this study are included in this published article.

## Author contributions

**Patricia Vega-Cuesta**: Conceptualization; Formal analysis; Validation; Investigation; Methodology; Writing-review and editing. **Ana López-Varea**: Investigation; Methodology. **Ana Ruiz-Gómez**: Formal analysis; Validation; Investigation; Visualization; Methodology; Supervision; Writing-review and editing. **Jose F. de Celis**: Formal analysis; Validation; Conceptualisation; Resources; Supervision; Writing-original draft; Writing-review and editing.

## Funding

This research was supported by Secretaría de Estado de Investigación, Desarrollo e Innovación, Grant/Award Number PID2022-141894OB-C21 to JFdC.

## Disclosure and competing interests statement

The authors declare no competing financial interests

## Acknowledgments

We thank the Developmental Studies Hybridoma Bank at Iowa University and Bloomington Stock Center for providing the tools necessary for this work and the support from the confocal microscopy CBMSO scientific service. We are very grateful to Diego Pulido, whose comments greatly improve this manuscript, and to Luis Alberto Baena for insights into the generation of Ras^Ko-Kin^ strains. The CBMSO enjoys institutional support from the Ramón Areces Foundation.

## Notes

### Competing Interest Statement

The authors have declared no competing interest.

